# A computational strategy for finding novel targets and therapeutic compounds for opioid dependence

**DOI:** 10.1101/359075

**Authors:** Xiaojun Wu, Siwei Xie, Lirong Wang, Peihao Fan, Songwei Ge, Xiang-Qun Xie, Wei Wu

## Abstract

Opioids are widely used for treating different types of pains, but overuse and abuse of prescription opioids have led to opioid epidemic in the United States. Besides analgesic effects, chronic use of opioid can also cause tolerance, dependence, and even addiction. Effective treatment of opioid addiction remains a big challenge today. Studies on addictive effects of opioids focus on striatum, a main component in the brain responsible for drug dependence and addiction. Some transcription regulators have been associated with opioid addiction, but relationship between analgesic effects of opioids and dependence behaviors mediated by them at the molecular level has not been thoroughly investigated. In this paper, we developed a new computational strategy that identifies novel targets and potential therapeutic molecular compounds for opioid dependence and addiction. We employed several statistical and machine learning techniques and identified differentially expressed genes over time which were associated with dependence-related behaviors after exposure to either morphine or heroin, as well as potential transcription regulators that regulate these genes, using time course gene expression data from mouse striatum. Moreover, our findings revealed that some of these dependence-associated genes and transcription regulators are known to play key roles in opioid-mediated analgesia and tolerance, suggesting that an intricate relationship between opioid-induce pain-related pathways and dependence may develop at an early stage during opioid exposure. Finally, we determined small compounds that can potentially target the dependence-associated genes and transcription regulators. These compounds may facilitate development of effective therapy for opioid dependence and addiction. We also built a database (http://daportals.org) for all opioid-induced dependence-associated genes and transcription regulators that we discovered, as well as the small compounds that target those genes and transcription regulators.

## Author summary

Opioids are widely used to treat pain in the clinics, however, overuse and abuse of prescription opioids in the United States cause opioid epidemic. There are no effective treatments for opioid addiction. Researchers have some understanding of the mechanism of opioid addiction at the molecular level, in relation to its pain-relieving effect in the brain, but there are still many issues to be addressed. We developed a computational strategy in an effort to find novel target genes and effective therapeutic treatment of opioid dependence and addiction. Using statistical and machine learning methods, we identified genes and transcription regulators that can serve as potential targets for treating opioid dependence and addiction. Our results revealed both known and novel genes and transcription regulators that were associated to dependence-related behavioral changes after opioid administration. Moreover, we found that many behavioral changes after opioid addiction are related to opioid effects on pain relief as well as immune and neuronal signaling. Following our analysis, we further determined small compounds that can potentially target dependence-associated genes and transcription regulators. We also built a database for the genes, transcription regulators, and small compounds, which is available at http://daportals.org.

## Introduction

Opioids such as morphine have long been used as mainstay therapy for treating different types of chronic severe pains such as cancer pain, noncancer-related pain, and neuropathic pain [1]. As a result of relaxation on restriction of prescription opioids for treating chronic noncancer pain and promotion of opioids in treatment by pharmaceutical industry, practitioners and many organizations, non-medical use and abuse of prescription opioids has been increasing rapidly in the United States [2], which in turn leads to opioid epidemic [3].

Despite that opioids have beneficial analgesic effects of alleviating acute and chronic pain, chronic opioid use can lead to adverse side effects including tolerance, hyperalgesia, withdrawal reactions, and even dependence [2, 4, 5]. In order to achieve optimal pain management, many studies have been carried out to elucidate mechanisms underlying both the beneficial as well as the adverse effects of opioids. Plenty of evidence has shown that crosstalks between neuronal signaling, immune responses and chemokines play significant roles in the pain pathways responsive to opioids such as morphine, and these signaling networks can lead to both behavioral and structural changes in the brain [6, 7].

Studies on effects of opioids to relieve pain mainly focus on brain regions such as periaqueductal gray, rostral ventromedial medulla, and dorsal root ganglia [6], while investigations of addictive effects of opioids mostly focus on striatum, since striatum is a main component of the reward system responsible for drug dependence and addiction. Striatum receives several types of neuronal inputs from prefrontal cortex (PFC), ventral tegmental area (VTA), and other areas of the brain [8], and over time, drug dependence and addiction can be reinforced [9]. At the molecular level, many transcription factors (TFs) have been associated with behaviors related to drug abuse and addiction. Some TFs known to play key roles in drug addiction include ΔFosB, cyclic AMP-responsive element binding protein (CREB), NF-κB, and MEF2 [7]. For instance, opioids can reduce Fos expression in the direct pathway striatal neurons [10]. Moreover, drugs of abuse can also alter gene transcription and induce addiction by epigenetic mechanisms [7].

Despite that different drugs of abuse often elicit similar behavioral responses in animals and humans, molecular mechanisms underlying addiction induced by different drugs can be distinctly different [10]. Opioids such as morphine and heroin increase dopamine level in nucleus accumbens, the main component of the ventral striatum, through activation of dopaminergic neurons in VTA [10]. It is popularly believed opioids inhibition of GABAergic neurons in VTA is another contributing factor of disinhibition of dopamine in VTA and increased rewarding effects in nucleus accumbens [11–16]. Opioids can also increase glutamate release in the nucleus accumbens, which results in changes in synaptic plasticity such as decreased dendritic branching and spine density [17].

Despite all these efforts, however, much remains to be known about molecular connections between the genes and pathways activated by opioids in the pain-related processes and those involved in opioid dependence and addiction. Elucidating such connections can not only shed light on the mechanisms which contribute to the opioid epidemic, but may also allow people to identify better candidate targets for therapeutic interventions to prevent opioid dependence and addiction during pain management.

In order to treat opioid addiction, opioid antagonists such as naltrexone have been used to treat opioid addiction for several decades. However, they have shown limited efficacy in relapse prevention [18, 19]. Recently, alternative approaches such as receptor-based therapeutic strategies have been proposed which aim to target receptors including G protein-coupled receptors (GPCRs) such as μ-, δ-, and κ-opioid receptors, chemokine receptors, as well as neuroimmune receptors such as Toll-like receptors 4 (TLR4) [20–22]. However, despite all the efforts and advances in understanding the mechanism of addiction in the past decades, they have not led to development of effective new anti-addiction agents [23]. Therefore, finding novel targets and strategies is needed for treating opioid dependence and addiction.

In this work, we developed a new computational strategy for identification of novel genome-wide targets and potential therapeutic treatments for opioid dependence. In particular, this strategy involves first detecting genes and pathways induced by either morphine or heroin which are associated with dependence, then identification of small compounds which can target the dependence-associated genes responsive to the opioids. Using this strategy, we identified morphine and heroin-induced genes and pathways associated with dependence-related behaviors such as physical dependence and psychological dependence, as well as transcription regulators which can potentially regulate these genes. Finally, we identified small compounds which can potentially target some of the dependence-related genes and transcription factors. A database for all the dependence associated genes and transcription regulators, along with small compounds for targeting those genes and transcription regulators, is available at http://daportals.org. Our findings can facilitate identification of novel candidate gene targets as well as potential therapeutic interventions for treating opioid dependence and addiction.

## Results

### Identification of genes and patterns induced by either morphine or heroin

In order to identify genes induced by either morphine or heroin over time, we applied a local regression method to gene expression microarray data collected from mice treated by each opioid at different time points [24]. Using this approach, we found that 423 genes were differentially expressed (DE) after morphine administration in mice, while 608 genes were differentially induced by heroin. Using *k*-means clustering, we identified 6 expression patterns among genes induced by each opioid (Figs 1 and 2), with genes upregulated and downregulated at 3 phases: Immediate-Early (IE), Middle (M), and Late (L), respectively. Compared to morphine, heroin induced twice as many genes in the IE and L phases, respectively (Figs 1 and 2), indicating that heroin elicits neurobiological responses in mice not only faster but also longer lasting than morphine.

**Fig 1:**
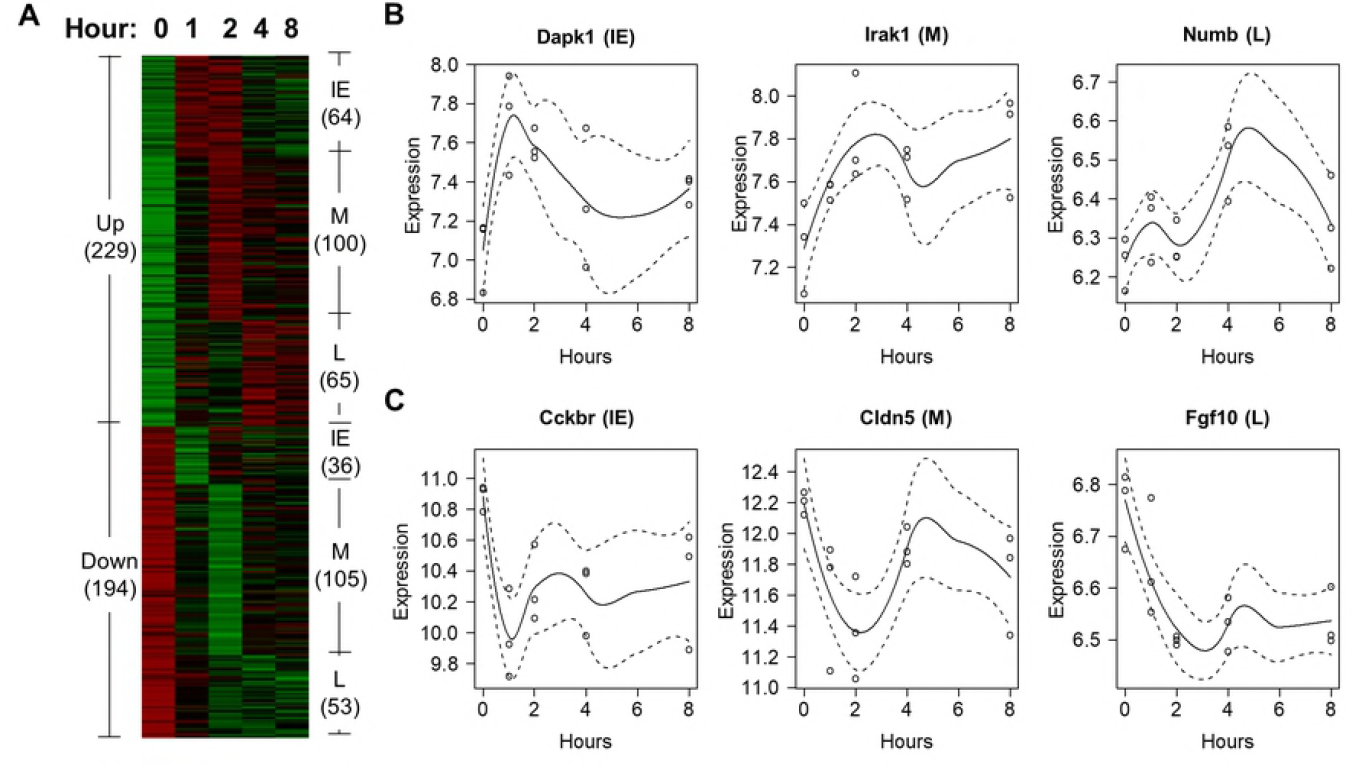
**Genes differentially expressed after morphine exposure in mouse striatum.** (A) Patterns of differentially expressed genes induced by morphine. (BC) These plots show six genes upregulated (B) and downregulated (C) by morphine in the IE, M, and L phase, respectively.

**Fig 2:**
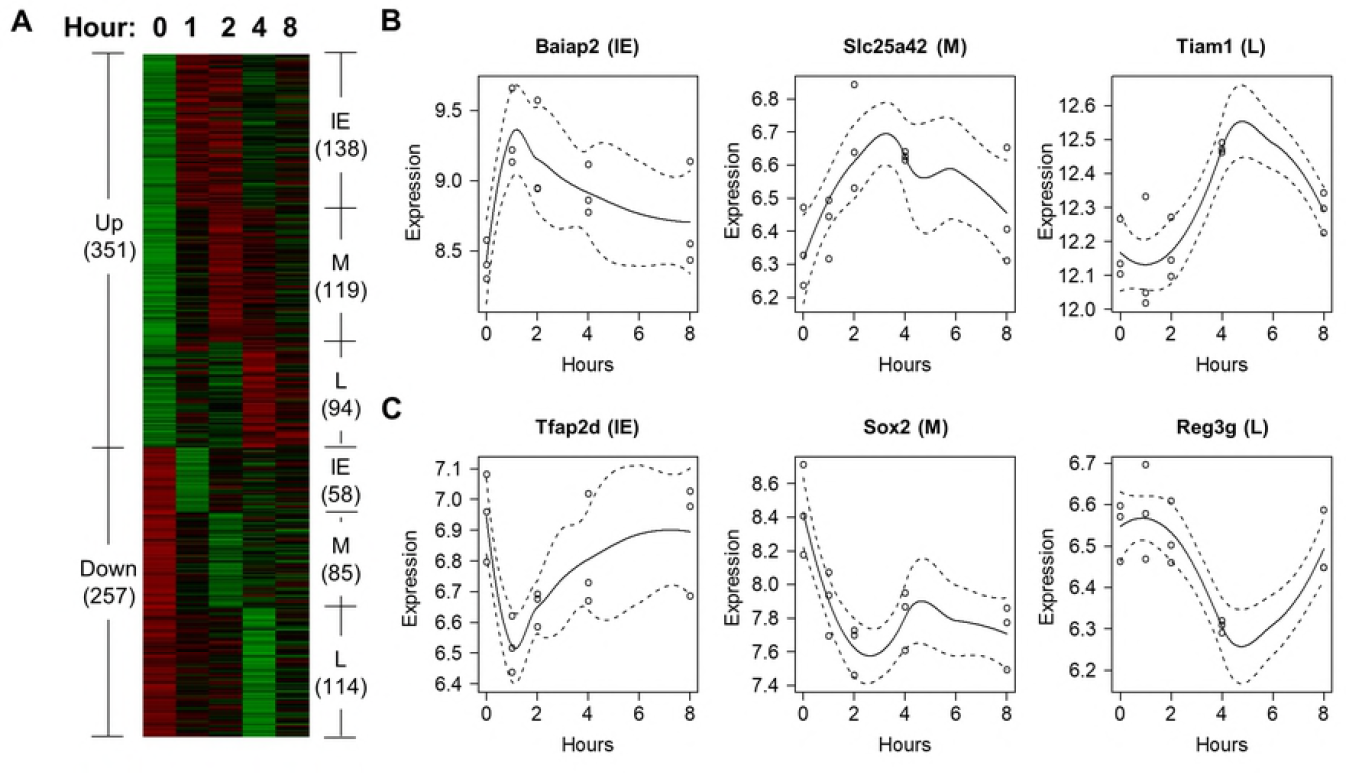
**Genes differentially expressed after heroin exposure in mouse striatum.** (A) Patterns of differentially expressed genes induced by heroin. (B-C) These plots show six genes upregulated (B) and downregulated (C) by heroin in the IE, M, and L phase, respectively.

### Enriched Gene Oncology (GO) and KEGG terms among differentially expressed genes (DEGs)

Our GO and KEGG analyses revealed that many DEGs induced by either morphine or heroin in the mouse striatum were involved in the immune and neuronal processes and pathways (Tables 1 and 2, S1 and S2 Tables). Many of these biological processes are previously known to play important roles in opioid-mediated pain pathways in brain regions such as dorsal root ganglia [6]. However, despite that the neuroimmune signaling processes induced by either morphine or heroin in mouse striatum share some similarities, it is also apparent that the two opioids elicit distinct neurobiological responses in the animals which we will detail below:

**Table 1:** **Significantly enriched biological processes and pathways induced by morphine which were involved in opioid-mediated pain pathways and corresponding literature support.** In the “Phase” column, Up-IE, Up-M, and Up-L represent upregulated in the IE, M, and L phase, respectively, while that Down-IE, Down-M, and Down-L represent downregulated in the IE, M, and L phase, respectively. Note: the full names of the behaviors can be found in S3 Table. HM* indicates that the DEG is induced by both heroin and morphine.

|  | GO/KEGG term | Phase | Effect (Literature support) | DEGs Associated with Dependence and Other Harmful Effects |
| --- | --- | --- | --- | --- |
| Immune system | Pattern recognition receptor signaling pathway | Up-M | Proinflammatory responses, tolerance [6, 22] | Irak1: phys dep; Ptafr: phys dep. |
|  | Activation of innate immune response |  | Proinflammatory responses, tolerance [6, 22] | Irak1: phys dep; Ptafr: phys dep. |
|  | Toll-like receptor signaling pathway |  | Allodynia and hyperalgesia [25, 26]; NF-κB activation [27-29] | Irak1: phys dep. |
|  | Response to xenobiotic stimulus | Up-L | Tolerance [30, 31] |  |
|  | Negative regulation of NF-κB TF activity |  | Analgesia<br>NF-κB activation [6] |  |
|  | MAPK signaling pathway | Down-M | Anti-inflammatory responses<br>Analgesia [32] |  |
|  | Humoral immune response |  |  |  |
|  | Induction of positive chemotaxis | Down-L | Nociceptive pathways [22] |  |
| Neuronal signaling | Cell projection morphogenesis | Up-L | Structural plasticity [33] | Numb: acute, dep, HCC, phys dep, phys harm |
|  | Calcium signaling pathway | Down-IE | Analgesia [22] |  |
|  | Synaptic transmission, glutamatergic | Down-M | Analgesia [34-36] | Grin1: phys dep, pleasure |
|  | Ensheatment of neurons |  | Nociceptive pathways [22] | Cldn5 (HM*): phys dep. |
|  | Synaptic transmission, GABAergic |  | Proinflammatory; tolerance [6, 37] |  |
|  | Glutamate receptor signaling pathway |  | Analgesia [6] | Cacng7: dep, phys dep, pleasure |
|  | Sensory organ development |  | Analgesia [38, 39] | Anp32b: phys dep. |
|  | Negative regulation of neuron apoptotic process |  | Neuronal apoptosis [6] | Grin1: phys dep, pleasure |
| Other key pathways | Protein dephosphorylation | Up-IE | Alleviating inflammatory hyperalgesia [6, 40] | Dusp12 (HM): phys dep. |
|  | positive regulation of autophagy |  | Production of proinflammatory cytokines, tolerance [6] | Plekhf1: psycho dep. |
|  | Regulation of programmed cell death |  | Production of proinflammatory cytokines, tolerance [6] | Dapk1 (HM): psycho dep; Plekhf1 (HM): psycho dep; Pim3 (HM): phys dep. |

**Table 2:** **Significantly enriched biological processes and pathways induced by heroin which were involved in opioid-mediated pain pathways and corresponding literature support.** All of the abbreviations used in this table can be found in the legend of Table 1.

|  | GO/KEGG term | Phase | Effect (Literature support) | DEGs Associated with Dependence and Other Harmful Effects |
| --- | --- | --- | --- | --- |
| Immune system | Response to biotic stimulus | Up-IE | Nociceptive pathways [6] | Ace: dep, phys dep; Baiap2: pleasure, psycho dep; Stab1 (HM): phys dep |
|  | MyD88-dependent toll-like receptor signaling pathway | Down-L | Anti-inflammatory effect; analgesia [30, 41] |  |
|  | T-helper 1 type immune response (Il4, Tlr6, Il27) |  |  |  |
| | Positive regulation of I-κB kinase/NF-κB signaling | | | |
|  | Inflammatory response |  |  |  |
|  | Regulation of MAP kinase activity |  |  |  |
|  | Microglial cell activation |  | Anti-inflammatory responses; analgesia [6] |  |
|  | Regulation of granulocyte chemotaxis |  | Nociceptive pathways [22] |  |
| Neuronal signaling | Cell morphogenesis involved in neuron differentiation | Up-IE | Nociceptive pathways [22] | Baiap2: pleasure, psycho dep |
|  | Regulation of nervous system development |  | Nociceptive pathways [22] | Ace: dep, phys dep; Ncs1: dep; Baiap2: pleasure, psycho dep |
|  | Regulation of excitatory postsynaptic membrane potential |  | Tolerance and hyperalgesia [22] | Sez6: dep |
|  | Potassium ion transport | Up-M | Proanalgesic effect [22] |  |
|  | Cation transmembrane transport |  | Tolerance and hyperalgesia [32] | Slc25a42: dep |
|  | Sodium ion transmembrane transport | Up-L | Allodynia and hyperalgesia [22] |  |
|  | Regulation of synapse organization |  | Analgesia [22] |  |
|  | Regulation of neuron death | Down-IE | Linked to anti-inflammatory response; analgesia [42] | Bag1: dep, pleasure; Tfap2d: dep, pleasure, psycho dep |
| Other key processes | Protein autophosphorylation | Up-IE | Hyperalgesia [22] | Dapk1 (HM): psycho dep; Pim3 (HM): phys dep |
|  | Negative regulation of receptor recycling | Up-L | Tolerance [22] | Pcsk9: pleasure, psycho dep |
|  | Response to cAMP | Down-M | Analgesia [6] |  |

### Immune and neuronal responses to morphine in mouse striatum

***Immune responses***: Most genes involved in the immune system were induced by morphine in the M phase. As shown in Table 1, some genes participating in anti-inflammatory processes were responsive to morphine, e.g., genes involved in negative regulation of NF-κB transcription factor activity were upregulated, and those involved in MAPK pathway, humoral immune response, and induction of positive chemotaxis were downregulated. Since anti-inflammatory pathways are known to play key roles in opioid-induced pain relief [6], these results are consistent with the fact that morphine induces analgesia effect in animals.

On the other hand, some genes participating in proinflammatory responses were also upregulated by morphine, including, e.g., genes involved in Toll-like receptor signaling pathway, and activation of innate immune response. Notably, previous evidence showed that proinflammatory responses play central roles which contribute to tolerance during chronic opioid exposure [7]. Genes involved in response to xenobiotic stimulus were also upregulated in the L phase, in line with the fact that these genes are essential in sensing that the cells are under the ‘insult’ of the drug.

Together, these results suggest that crosstalks between genes involved in proinflammatory and anti-inflammatory pathways have already initiated in mouse striatum during short-term morphine administration.

***Neuronal responses***: Our results also showed that genes participating in neuronal responses were induced by morphine (Table 1), which is not surprising, given known neurological effects of morphine in the brain. For example, positive regulation of autophagy and programmed cell death were upregulated in the IE phase, and also genes involved in glutamatergic synaptic transmission, glutamate receptor signaling pathway, and sensory organ development were all downregulated in the M phase. Notably, all these neuronal signaling events have been implicated in the analgesia induced by morphine (Table 1), again supporting the notion that morphine can induce analgesia in the treated mice. Also, genes involved in cell projection morphogenesis were upregulated in the L phase, in line with the evidence that morphine can induce structural changes in mice during chronic exposure [33].

Together, our results suggest that even during short-term morphine exposure, complex crosstalks between genes involved in proinflammatory and anti-inflammatory pathways, neuronal signaling, and the chemokine system have already initiated in mouse striatum, which can contribute to analgesia and/or tolerance effects if exposure of the drug lasts longer.

### Immune and neuronal responses to heroin in mouse striatum

***Immune responses***: Our results showed that distinct from morphine, many immune genes induced by heroin were downregulated in the L phase (Table 2), which included MyD88-dependent Toll-like receptor signaling pathway, microglial cell activation, T-helper 1 type immune response (Il4, Tlr6, Il27), positive regulation of I-κB kinase/NF-κB signaling, regulation of granulocyte chemotaxis, inflammatory response, and regulation of MAP kinase activity. Notably, all these pathways are known to be active in proinflammatory responses during chronic opioid exposure [6]. Therefore, downregulation of these pathways indicates that heroin induces strong anti-inflammatory response and thus elicits strong analgesic effects in the mice.

***Neuronal responses***: Genes involved in neuronal activities, such as regulation of nervous system development and excitatory postsynaptic membrane potential (IE phase) and cation transmembrane transport (M phase) were upregulated by heroin, whereas genes involved in regulation of neuron death were downregulated (IE phase) (Table 2). These results also agree with previous findings that neuronal responses play active roles in opioid-related pain process [6].

Also, we noticed that genes involved in cell morphogenesis involved in neuron differentiation, positive regulation of axonogenesis, and regulation of synapse organization were upregulated among IE and L phases after heroin exposure. These results are supported by the previous evidence that drugs of abuse can induce changes in structural plasticity in animals during chronic exposure [43].

***Other key biological responses***: Our results also showed that heroin induced genes participating in other key biological processes involved in pain-related pathways, e.g., genes involved in protein autophosphorylation are upregulated in the IE phase. Since protein phosphorylation is known to play key roles in desensitization and implicated in opioid-induced hyperalgesia [22], our results suggest that these genes contribute to dependence induced by chronic use of heroin; this speculation is confirmed by our association analysis described below.

### Association of morphine- and heroin-induced DEGs with harmful effects of drugs of abuse

In order to find out whether DEGs induced by either morphine or heroin are associated with various harmful effects linked to drugs of abuse, we conducted the association analysis between expression levels of morphine- or heroin-induced DEGs and twelve DA-related harmful effects including dependence, physical dependence, psychological dependence, pleasure, physical harm, social harm, health care cost, and conditioned place preference (See S3 Table and Methods for details).

Using this approach, we detected 44 morphine-induced DEGs and 61 heroin-induced DEGs significantly associated with dependence-related behaviors at the nominal level of significance (p<0.05) (S4 and S5 Tables). Among these dependence-associated DEGs, 9 were induced by both morphine and heroin, of which 6 were induced in the IE phase. These results can be searched in our database at http://daportals.org.

Next, we investigated whether the dependence-associated DEGs induced by either morphine or heroin were involved in the pain-related neuroimmune pathways mediated by opioids. As shown in Tables 1 and 2, we found that a significant number of the dependence-associated DEGs induced by the opioids were involved in the pain-related neuroimmune pathways. Moreover, some of these genes could be induced by both morphine and heroin, e.g., Dapk1, Plekhf1, Pim3, and Dusp12 were upregulated by both morphine and heroin in the IE phase, and were associated with psychological dependence and physical dependence, respectively.

### Detection of potential transcription regulators that regulate the opioid-induced dependence-associated DEGs

In order to find out whether the dependence-associated DEGs induced by each opioid were co-regulated by any TFs or epigenetic factors, we performed the TF and epigenetic factor binding site enrichment test using the ENCODE ChIP-Seq significance tool [44]. Our analysis showed that 17 transcription regulators potentially modulated morphine-responsive dependence-associated DEGs (S6 Table), while that 12 transcription regulators modulated heroin-responsive dependence-associated DEGs (S7 Table). More details about our results are described below.

### Transcription regulators detected after morphine exposure

***Known TFs associated with dependence and addiction***: More than half of the TFs we detected regulating dependence-associated DEGs after morphine exposure are previously known to play important roles in drug dependence and addiction. For example, we found that MEF2A upregulated four DEGs which were associated with physical dependence in the M phase, while that MEF2C upregulated one DEG associated with dependence in the IE phase after morphine exposure, agreeing with the evidence that MEF2 is crucial in inducing behavioral changes after exposure to drugs of abuse [7].

***Novel TFs induced by morphine***: Our results also showed a few novel TFs activated by morphine. In order to quantify the magnitude of the effects the detected transcription regulators on the dependence-related behaviors induced by the opioids, we developed a scoring metric, called dependence score, based on the total fold changes of the dependence-associated DEGs co-regulated by each regulator (see details in Methods). Using this approach, we found that E2f6, which potentially regulated 13 DEGs associated with physical dependence after morphine exposure, had the highest dependence score of 17.93. Also, we detected ZBTB33 and ZKSCAN1 associated with physical dependence having high dependence scores (11 and 6.8, respectively) after morphine exposure. In particular, ZBTB33 encodes a transcriptional regulator Kaiso which can promote histone deacetylation and decrease expression levels of its target genes [45], consistent with our results showing that ZBTB33 downregulates 8 genes in the M phase. Also, Zkscan1 encodes a member of the Kruppel C2H2-type zinc-finger family of proteins, which has been implicated in regulating the expression of GABA type-A receptors in the brain [46].

***Epigenetic factors***: Our results showed that about half (8 out of 17) of the detected transcription regulators after morphine exposure were epigenetic factors (S6 Table). Literature search (S8 Table) suggests that these factors including HDAC8 (encoding histone deacetylase 8) and HDAC6 (encoding histone deacetylase 6) play important roles in histone acetylation, histone methylation, DNA methylation, and chromatin remodeling [45, 47–52], in line with the previous evidence that all these epigenetic events have been implicated in the neurobiological responses to drugs of abuse in the brain [7]. Notably, among all the epigenetic factors, SAP30 which encodes a component of the histone deacetylase complex had the highest dependence score of 15.49 and can potentially co-regulate 11 morphine-responsive DEGs associated with physical dependence.

### Transcription regulators detected after heroin exposure

***Known TFs associated with dependence and addiction***: DEGs upregulated by EGR1 and CREB1 were associated with psychological dependence in the IE phase after administration of heroin, consistent with the fact that both EGR1 and CREB are key TFs which regulate genes involved in dependence-related behavioral responses during exposure of opioids as well as other drugs of abuse [5, 7]. Moreover, our results showed that Polr2a (encoding the largest subunit of RNA polymerase II), EGR1, and CREB1 were associated with social and physical harm with the highest scores (> 20), suggesting that these TFs play key roles in the biological mechanism that underlies the higher social and physical harm caused by heroin compared to other drugs of abuse [4].

***Novel TFs induced by heroin***: We found that E2F6 (which showed the highest dependence score in morphine) also had the highest dependence score of 6.52, potentially regulating four DEGs associated with psychological dependence after exposure to heroin.

***Epigenetic regulators***: Similar to morphine, more than 40% of the detected transcription regulators were epigenetic factors after heroin exposure (S7 Table). All these factors have been known to play major roles in histone and DNA methylation, and chromatin remodeling methylation [50–53]. Notably, 3 epigenetic factors CTCF, EZH2, and SUZ12 were identified during both morphine and heroin exposure.

Taken together, our results suggest that distinct transcriptional regulatory mechanisms are responsive to exposure of morphine and heroin in mouse striatum, and that epigenetic regulation plays major roles after exposure to both opioids. Further investigation is needed to elucidate the roles of the novel TFs (e.g., E2F6) with high dependence scores after exposure of either morphine or heroin.

### Finding small compounds which can target the DEGs and the dependence-associated transcription regulators induced by the opioids

To facilitate development of therapeutic interventions for treating morphine or heroin dependence, we developed a strategy which allowed us to identify small compounds that can target the opioid-induced dependence-associated DEGs and their potential transcription regulators (see details in Methods). Our results are shown in S9 and S10 Tables for morphine and heroin, respectively. Among all the small compounds we identified, we found that Calmidazolium could target 8 morphine-induced dependence-associated DEGs (S9A Table), and that Securinine could target E2F6 which had the highest dependence score after morphine exposure (Table 3). Also, we identified that Phenacetin and Buspirone could target 11 and 4 heroin-induced dependence-associated DEGs, respectively (Table 4, S1 Fig), and that Meclofenoxate could target E2F6 which had the highest dependence score also after heroin exposure (S10B Table).

**Table 3:** **Small compounds negatively correlated with the potential transcription regulators after morphine administration.** All of the abbreviations used in this table can be found in the legend of Table 1.

| Compound | Transcription Regulators | Fold Change | Phase | Associated Harmful Effects |
| --- | --- | --- | --- | --- |
| Benserazide | TAF1 | 2.9 | Up-IE | Dep, pleasure |
|  | MEF2A | 5.34 | Up-M | Phys dep |
|  | ZKSCAN1 | -6.81 | Down-M | Phys dep |
| Piperacetazine | SUZ12 | 1.2 | Up-IE | Dep |
|  | ZKSCAN1 | -6.81 | Down-M | Phys dep |
| Securinine | MEF2C | 1.7 | Up-IE | Dep |
|  | BRF2 | 1.39 | Up-M | Phys dep |
|  | E2F6 | -17.93 | Down-M | Phys dep |
| Isocorydine | MEF2A | 5.34 | Up-M | Phys dep |
| Gabapentin | CTCF | 2.97 | Up-IE | Chronic, dep, phys harm |
|  | MEF2A | 5.34 | Up-M | Phys dep |
|  | HDAC6 | -4.84 | Down-M | Acute, phys dep |

**Table 4:**
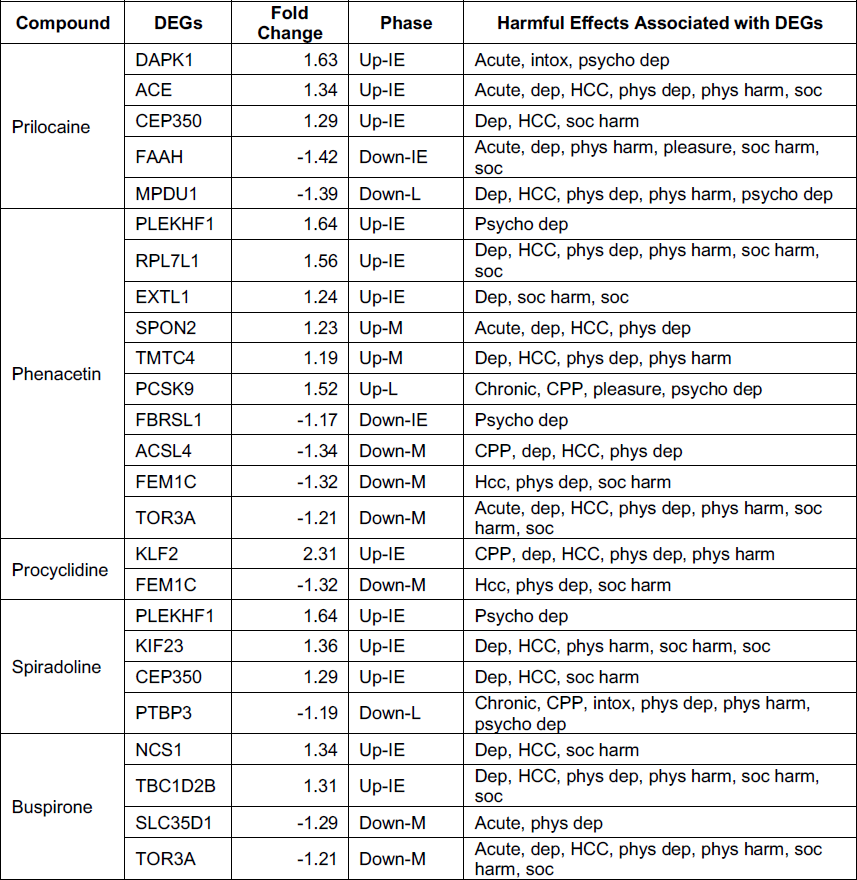
**Small compounds negatively correlated with dependence-associated DEGs after heroin administration.** All of the abbreviations used in this table can be found in the legend of Table 1.

## Discussion

Many studies have been conducted which intend to delineate pain-related pathways induced by opioids. However, much remains to be known about the molecular connection between these opioid-mediated pain pathways and those playing key roles in drug dependence and addiction. Dissecting these pathways can facilitate identification of candidate targets for developing effective therapeutic interventions which ideally can target opioid tolerance and dependence while preserving opioid analgesic effect.

In this study, we developed a computational strategy to identify candidate dependence-associated DEGs induced by either morphine or heroin, as well as to find small compounds which could target these genes for treating dependence of the opioids. Using this strategy, we analyzed a time-course gene expression microarray data set generated previously to investigate gene expression patterns responsive to various drugs of abuse in mouse striatum [24]. In particular, we first employed a local regression technique to detect genes differentially expressed over 8 hours of time in mouse striatum after either morphine or heroin exposure. Then, we performed correlation analysis to identify morphine or heroin-induced DEGs which were associated with twelve harmful effects including dependence commonly linked to drugs of abuse. Furthermore, we detected potential transcription regulators including TFs and epigenetic factors that regulated the dependence-associated DEGs using an ENCODE enrichment tool. Finally, to facilitate the identification of candidate targets and development of effective therapy for morphine and heroin-induced dependence, we identified small compounds which could potentially target against some of the detected dependence-associated DEGs and transcription regulators.

Using the approach described above, we found that a significant number of the DEGs responsive to either morphine or heroin in mouse striatum were involved in the neuroimmune signaling pathways, which are typically activated in the pain-related pathways during chronic opioid use previously identified in other brain areas including periaqueductal gray, rostral ventromedial medulla, and dorsal root ganglia [6]. Using correlation analysis, we found that a considerable portion of the pain pathway-related DEGs, previously known to play active roles in opioid analgesia, tolerance, hyperalgesia, and allodynia, were associated with the harmful effects (such as dependence) linked to morphine and heroin as well as many other drugs of abuse, e.g., Irak1 (encoding interleukin-1 receptor-associated kinase 1) in the enriched Toll-like receptor signaling pathway (Table 1) induced by morphine was correlated with physical dependence at a nominal level of significance (p < 0.05). Toll-like receptor signaling pathway has been known to play crucial roles in proinflammatory signaling and tolerance to opioid analgesia. We also noticed that some dependence-associated DEGs could be induced by both morphine and heroin, e.g., among DEGs upregulated in the IE phase, Dapk1 and Plekhf1 were correlated with psychological dependence, while Dusp12 and Pim3 were associated with physical dependence after exposure to both morphine and heroin. It is unclear what roles these genes (induced by both opioids) play when mice were first exposed to morphine, then switched to heroin later on, a scenario commonly seen among human drug abusers.

Despite the similarities in gene expression responses induced by both morphine and heroin in mouse striatum, differences between the two opioids are also obvious. For example, a large number of the DEGs involved in immune signaling were downregulated in the L phase after heroin exposure, as opposed to morphine exposure, suggesting that heroin elicited strong anti-inflammatory responses in the L phase and thus induced acute analgesic effects in the mice.

Among detected transcription regulators that potentially regulate the dependence-associated DEGs, MEF2A induced by morphine as well as EGR1 and CREB1 by heroin are known to play crucial roles in drug addiction. We also found that more than 40% of the detected transcription regulators are epigenetic factors after both morphine and heroin exposure, including HDAC8 and HDAC6 activated by morphine, which supports the previous notion that epigenetic factors are important for addiction. Furthermore, using the dependence score, a metric we developed for measuring the extent the detected transcription regulators affect the dependence-associated DEGs, we found that E2F6 has the highest dependence scores after exposure of both morphine and heroin.

In summary, our work here intent to elucidate molecular connections between the analgesic and tolerance-related pain pathways and harmful side effects of opioid use during pain treatment. Despite the general belief that morphine is safe for managing patients with pain, our results suggest that morphine may induce tolerance to analgesia and dependence on the drug in the patients in the very early stage, which may increase the possibility of the same patients to abuse heroin thereafter, since heroin may further induce acute analgesic effects as suggested by our results. Moreover, because heroin can cause both structural and behavioral changes among patients, abusing heroin after morphine may lead to more potent dependence on the drugs among the patients.

Furthermore, we found several small compounds which could potentially target some of the dependence-associated DEGs and the detected transcription regulators induced by the opioids. In particular, we identified Securinine and Meclofenoxate which could target E2F6 in humans after exposure to morphine and heroin, respectively. These compounds can facilitate future development of effective therapeutic interventions which can target the adverse side effects of morphine and heroin, while preserving their analgesic effects.

The limitations of our work include the following. The gene expression microarray data we analyzed spanned only 8 hours after administration of morphine and heroin, which limited our ability to discover chronic effects of the drugs. In the future study, we intend to employ the same strategy to investigate long-term effects of morphine and heroin, and to compare them with the acute effects we discovered in this work. Also, biological validation is needed to verify our findings here.

Despite the limitations, we found that our results agree well with the previously known evidence about drug abuse and addiction, suggesting our findings are valid and worth further in-depth investigation. Moreover, our work provides insight into the molecular connections between the opioid-induced pain-related pathways and the adverse harmful effects associated with morphine and heroin. Understanding such connections may facilitate development of effective therapies which allow people to target dependence-associated genes and transcription regulators at an early stage of opioid use while preserving analgesic effects of opioids.

## Methods

### Dataset

The gene expression microarray data set we analyzed in this work was obtained from the NCBI Gene Expression Omnibus (GEO) database under the accession number [GEO:GSE15774]. This data set was generated from a previous work described in [24], in which, gene expression alterations in mouse striatum were investigated after the mice were treated by various drugs of abuse, including morphine, heroin, methamphetamine, cocaine, nicotine and alcohol. Detailed description of the data set can be found in [24]. Briefly, after a single dose of drug administration, gene expression was obtained from the mouse striatum at 1, 2, 4, 8 hours afterwards. Meanwhile, samples from saline- and naïve-treated control group were collected at 0, 1, 2, 4, 8 hours as controls. There were three biological replicates for each drug group and each time point.

### Identification of genes differentially expressed over time in mouse striatum after exposure of either morphine or heroin

In order to identify genes differentially expressed over time in mouse striatum after administration of either morphine or heroin, we employed a local regression smoothing technique [54] to estimate the smoothed time course gene expression data for each opioid. The detailed description of the strategy can be found in [55]. For each opioid, expression values of each gene (i.e., transcript) were available for 1, 2, 4, and 8 hours, and expression values for time point 0 for the corresponding genes from the naïve group were used to represent the control time point (i.e., 0 hour) for each opioid. In particular, expression values for each gene over different time points were first fitted using a local polynomial quadratic (degree = 2) model with the bandwidth optimally estimated using a leave-one-out cross validation procedure [54]. To determine whether a gene is differentially expressed over time with respect to the control time point, we calculated the simultaneous 95% confidence intervals for the fitted (or expected) intensity values using a method due to Sun and Loader [56]. The p-values were adjusted using the Bonferroni correction to account for multiple hypothesis testing. We determined a gene as differentially expressed if its expression value at any time point *T* relative to the control time point satisfied: 1) adjusted p-value < 0.05, and 2) fold change ≥ 1.2.

### Identification of temporal patterns for DEGs induced by either morphine or heroin using cluster analysis

In order to identify temporal patterns for DEGs responsive to either morphine or heroin exposure, we applied a k-means clustering algorithm proposed by Hartigan and Wong [57] to the temporal expression values of the DEGs. The Euclidean distance was used to measure dissimilarities between different genes. A thousand iterations were performed to find an optimal partition of *K* clusters where *K* is preassigned. To determine an optimal number of the clusters for the DEGs, we employed the average silhouette width (ASW) as described in [58], and when *K* = 6, ASW is the largest for the DEGs induced by both morphine and heroin.

### GO and KEGG pathway enrichment analysis

We performed GO and KEGG pathway enrichment analysis to identify biological processes and pathways that were overrepresented among DEGs in each cluster after exposure of either morphine or heroin. We performed the enrichment analysis with the GOstats R software package [59], which finds enriched functional groups using the hypergeometric test with the aid of the functional terms in the GO and KEGG databases. GO terms and KEGG pathways were considered as significantly enriched if their p-values < 0.05.

### Association of morphine- and heroin-induced DEGs with harmful effects of drugs of abuse

Our association analysis aimed to determine whether morphine- and heroin-induced DEGs were associated with any harmful effects of drugs of abuse. The scores which assessed the magnitudes of the twelve harmful effects of various drugs of abuse were taken from Nutt et. al. [4] and can be found in S3 Table. In particular, the scores for the harmful effects encompassing three categories, including physical harm (overall, acute, chronic), dependence (pleasure, psychological, physical), social harm (overall, health-care costs), and conditioned place preference for the drugs including morphine, heroin, cocaine, methamphetamine, ethanol, and nicotine were used to calculate the association of each harmful effect and the DEGs induced by either morphine or heroin. Specifically, let *S_i_* denote a vector of the scores for harmful effect *i* corresponding to drugs *D*, where *D* = [morphine, heroin, cocaine, methamphetamine, ethanol, and nicotine]. Let *G_j_* denote a vector of expression values of gene *G* corresponding to drugs *D* at time point *j*; *G_j_* is a DEG induced by either morphine or heroin at time point *j*, but the gene is not required to be differentially induced by the other drugs at the same time point. The expression values of gene *G* for the drugs including cocaine, methamphetamine, ethanol, and nicotine were estimated for each drug by using the same local regression smoothing techniques as described above for morphine and heroin. Finally, we calculated the correlation between *G_j_* and *S_i_* using both the Pearson correlation and a quadratic polynomial regression; if the resulting p-value from any of the methods was less than 0.05, *G_j_* was considered as significantly associated with *S_i_*.

### Identification of human transcription and epigenetic factors that potentially regulate the dependence-associated DEGs induced by each opioid

To identify potential transcription and epigenetic factors that regulate dependence-associated DEGs responsive to each opioid in mouse striatum, we employed the ENCODE ChIP-Seq significance tool [44] to identify human TFs and epigenetic factors whose binding sites were significantly enriched among the DEGs associated with dependence (i.e., dependence, psychological, and/or physical dependence) in each of the six identified clusters after either morphine or heroin exposure. The ENCODE ChIP-Seq significance tool calculates enrichment scores of the transcription regulators using the hypergeometric test, and the resulting p-values were corrected by an FDR procedure to account for multiple hypothesis testing. A 1000-base pair (bp) window upstream of the transcription start site (TSS) and downstream of the transcription termination site (TTS) were considered for each DEG. A TF or an epigenetic factor was considered as significantly enriched if its FDR p-value < 0.05.

Furthermore, to facilitate ranking the significantly enriched TFs and epigenetic factors in terms of their impact on dependence, we developed a scoring metric called the ‘dependence score’ as follows. For each transcription regulator *R* activated in a certain phase *P*, we assume that *R* regulates a number *N* of morphine or heroin-induced DEGs associated with a dependence-related harmful effect (such as dependence, psychological, or physical dependence) within phase *P*. As shown in Figs 1 and 2, the IE, M, and L phases correspond to 0-2 hours, 2-4 hours, and 4-8 hours after exposure to either morphine or heroin. The ‘dependence score’ for *R* in phase *P* was then defined as 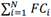 where *FC_i_* represents the maximum absolute fold change of the expression values of a DEG G*i* within phase P, relative to that of the control time point, and *G_i_* is regulated by R. The higher the dependence score, the more impact the transcription regulator *R* can have on dependence.

Using a similar concept as the dependence score, we also assigned association scores to transcription regulator *R* if the DEGs it regulates were also associated with other harmful effects of drugs (as shown in S3 Table). Specifically, an association score between a transcription regulator *R* and a harmful effect *H* was defined as the sum of the maximum absolute fold change of the DEGs regulated by *R* in phase *P* that were associated with *H*.

### Finding small molecular compounds to target the opioid-induced dependence-associated DEGs and the transcription regulators

Gene expression patterns in cells can change during treatment by small-molecule drugs or compounds [60]. If a small compound has an opposite effect on transcription than opioids, the small compound has potential to reverse the gene signature induced by opioids and hence the subsequent harmful effects caused by opioids. With the availability of gene expression profiles of small molecular compounds, we were able to compare them with those of morphine and heroin, and identify small compounds with the potential for treating dependence and addiction induced by each opioid.

Specifically, we employed the following two-step strategy to find the small compounds:

### Finding DEGs induced by small compounds

First, a commercial Illumina BaseSpace (former Nextbio™) software (Santa Clara, CA, USA, http://www.nextbio.com) were used to obtain the DEGs induced by small compounds in cells. In BaseSpace, most of the raw gene expression datasets involving perturbations by small compounds were obtained from the Gene Expression Omnibus (GEO) database (http://www.ncbi.nlm.nih.gov/geo/). Only genes with the p-values < 0.05 and absolute fold changes >1.2 were considered as DEGs induced by small compounds. To identify top compounds that have gene expression profiles most correlated with the dependence-associated DEG or the transcription regulators induced by morphine or heroin, we searched the gene expression profiles of small compounds stored in BaseSpace through BaseSpace integrated Pharmaco Atlas search. Then, the correlation between the DEGs induced by small compounds and the opioid-induced DEGs or transcription regulators was calculated as described below.

### Calculation of the correlation between the DEGs induced by small compounds and the opioid-induced DEGs or transcription regulators

For each opioid-induced dependence-associated DEG, we used its maximum absolute fold change over the measured time points (i.e., 1, 2, 4, and 8 hours) (which was defined as the ratio of the highest absolute expression value of the gene relative to that at the control time point) to represent its fold change. For each transcription regulator, we used its dependence score to represent its fold change value.

The correlation between the DEGs induced by each small compound and the opioid-induced dependence-associated DEGs or transcription regulators was calculated using the BaseSpace software. This software provided a modified form of the rank-based enrichment statistics to compare the two sets of the DEGs [61, 62]. BaseSpace pre-processed gene expression data with biomedical ontologies to enable comparison among heterogeneous datasets from different species. It also used meta-analyses to provide consistent predictions from multiple instances of similar perturbations, e.g., genes expression profiles from different cell lines induced by the same compounds [63]. All analyses using the BaseSpace software were performed with the default parameters.

S1 Fig shows an example of the significant negative correlation between the (61) heroin-induced dependence-associated DEGs and the buspirone-induced DEGs (p-value = 0.0277). Four genes were regulated by both heroin and buspirone, but in opposite directions.

## Supporting information

**S1 Fig: Significant negative correlation between the 61 heroin-induced dependence-associated DEGs and the buspirone-induced DEGs (p-value = 0.0277).** Four genes were regulated by both heroin and buspirone, but in opposite directions.

**S1 Table: Significantly enriched GO terms and KEGG pathways induced by morphine in different phases and their association with harmful effects of drugs.** (A) Significantly enriched GO terms induced by morphine. (B) Significantly enriched KEGG pathways induced by morphine.

**S2 Table: Significantly enriched GO terms and KEGG pathways induced by heroin in different phases and their association with harmful effects of drugs.** (A) Significantly enriched GO terms induced by heroin. (B) Significantly enriched KEGG pathways induced by heroin.

**S3 Table: The scores of the harmful effects associated with drugs of abuse (adapted from** ***Nutt, et. al., Lancet 2007; 369: 1047–53***).

**S4 Table: 44 dependence-associated DEGs induced after morphine exposure.**

**S5 Table: 61 dependence-associated DEGs induced after heroin exposure.**

**S6 Table: Significantly enriched transcription regulators and associated harmful effects after morphine exposure.**

**S7 Table: Significantly enriched transcription regulators and associated harmful effects after heroin exposure.**

**S8 Table: Supporting evidence of involvement of transcription regulators in drug dependence and addiction from literature.**

**S9 Table: Small compounds negatively correlated with dependence-associated DEGs and the potential transcription regulators after morphine administration.** (A) Small compounds negatively correlated with dependence-associated DEGs induced by morphine. (B) Small compounds negatively correlated with the transcription regulators induced by morphine.

**S10 Table: Small compounds negatively correlated with dependence-associated DEGs and the potential transcription regulators after heroin administration.** (A) Small compounds negatively correlated with dependence-associated DEGs induced by heroin. (B) Small compounds negatively correlated with the transcription regulators induced by heroin.

## References

1. Pergolizzi J, Boger RH, Budd K, Dahan A, Erdine S, Hans G, et al. Opioids and the management of chronic severe pain in the elderly: consensus statement of an International Expert Panel with focus on the six clinically most often used World Health Organization Step III opioids (buprenorphine, fentanyl, hydromorphone, methadone, morphine, oxycodone). Pain Pract. 2008;8(4):287–313. Epub 2008/05/28. doi: 10.1111/j.1533-2500.2008.00204.x. PubMed PMID: 18503626.

2. Brady KT, McCauley JL, Back SE. Prescription Opioid Misuse, Abuse, and Treatment in the United States: An Update. Am J Psychiatry. 2016;173(1):18–26. Epub 2015/09/05. doi: 10.1176/appi.ajp.2015.15020262. PubMed PMID: 26337039; PubMed Central PMCID: PMCPMC4782928.

3. Manchikanti L, Helm S, 2nd, Fellows B, Janata JW, Pampati V, Grider JS, et al. Opioid epidemic in the United States. Pain Physician. 2012;15(3 Suppl):ES9-38. Epub 2012/07/20. PubMed PMID: 22786464.

4. Nutt D, King LA, Saulsbury W, Blakemore C. Development of a rational scale to assess the harm of drugs of potential misuse. Lancet. 2007;369(9566):1047–53. Epub 2007/03/27. doi: 10.1016/S0140-6736(07)60464-4. PubMed PMID: 17382831.

5. Ron D, Jurd R. The “ups and downs” of signaling cascades in addiction. Sci STKE. 2005; 2005(309):re14. Epub 2005/11/10. doi: 10.1126/stke.3092005re14. PubMed PMID: 16278489.

6. Hutchinson MR, Shavit Y, Grace PM, Rice KC, Maier SF, Watkins LR. Exploring the neuroimmunopharmacology of opioids: an integrative review of mechanisms of central immune signaling and their implications for opioid analgesia. Pharmacol Rev. 2011;63(3):772–810. Epub 2011/07/15. doi: 10.1124/pr.110.004135. PubMed PMID: 21752874; PubMed Central PMCID: PMCPMC3141878.

7. Robison AJ, Nestler EJ. Transcriptional and epigenetic mechanisms of addiction. Nat Rev Neurosci. 2011;12(11):623–37. Epub 2011/10/13. doi: 10.1038/nrn3111. PubMed PMID: 21989194; PubMed Central PMCID: PMCPMC3272277.

8. Yager LM, Garcia AF, Wunsch AM, Ferguson SM. The ins and outs of the striatum: role in drug addiction. Neuroscience. 2015;301:529–41. Epub 2015/06/28. doi: 10.1016/j.neuroscience.2015.06.033. PubMed PMID: 26116518; PubMed Central PMCID: PMCPMC4523218.

9. Everitt BJ, Robbins TW. Neural systems of reinforcement for drug addiction: from actions to habits to compulsion. Nat Neurosci. 2005;8(11):1481–9. Epub 2005/10/28. doi: 10.1038/nn1579. PubMed PMID: 16251991.

10. Badiani A, Belin D, Epstein D, Calu D, Shaham Y. Opiate versus psychostimulant addiction: the differences do matter. Nat Rev Neurosci. 2011;12(11):685–700. Epub 2011/10/06. doi: 10.1038/nrn3104. PubMed PMID: 21971065; PubMed Central PMCID: PMCPMC3721140.

11. Steidl S, Wasserman DI, Blaha CD, Yeomans JS. Opioid-induced rewards, locomotion, and dopamine activation: A proposed model for control by mesopontine and rostromedial tegmental neurons. Neurosci Biobehav Rev. 2017;83:72–82. Epub 2017/09/28. doi: 10.1016/j.neubiorev.2017.09.022. PubMed PMID: 28951251; PubMed Central PMCID: PMCPMC5730464.

12. Gysling K, Wang RY. Morphine-induced activation of A10 dopamine neurons in the rat. aBrain Res. 1983;277(1):119–27. Epub 1983/10/24. PubMed PMID: 6315137.

13. Johnson SW, North RA. Opioids excite dopamine neurons by hyperpolarization of local interneurons. J Neurosci. 1992;12(2):483–8. Epub 1992/02/01. PubMed PMID: 1346804.

14. Klitenick MA, DeWitte P, Kalivas PW. Regulation of somatodendritic dopamine release in the ventral tegmental area by opioids and GABA: an in vivo microdialysis study. J Neurosci. 1992;12(7):2623–32. Epub 1992/07/01. PubMed PMID: 1319478.

15. Margolis EB, Hjelmstad GO, Fujita W, Fields HL. Direct bidirectional muopioid control of midbrain dopamine neurons. J Neurosci. 2014;34(44):14707-16. Epub 2014/10/31. doi: 10.1523/JNEUROSCI.2144-14.2014. PubMed PMID: 25355223; PubMed Central PMCID: PMCPMC4212068.

16. Matthews RT, German DC. Electrophysiological evidence for excitation of rat ventral tegmental area dopamine neurons by morphine. Neuroscience. 1984;11(3):617–25. Epub 1984/03/01. PubMed PMID: 6717805.

17. Russo SJ, Dietz DM, Dumitriu D, Morrison JH, Malenka RC, Nestler EJ. The addicted synapse: mechanisms of synaptic and structural plasticity in nucleus accumbens. Trends Neurosci. 2010;33(6):267–76. Epub 2010/03/09. doi: 10.1016/j.tins.2010.02.002. PubMed PMID: 20207024; PubMed Central PMCID: PMCPMC2891948.

18. Kosten TA, Kosten TR. Pharmacological blocking agents for treating substance abuse. J Nerv Ment Dis. 1991;179(10):583–92. Epub 1991/10/01. PubMed PMID: 1919542.

19. Soyka M, Mutschler J. Treatment-refractory substance use disorder: Focus on alcohol, opioids, and cocaine. Prog Neuropsychopharmacol Biol Psychiatry. 2016;70:148–61. Epub 2015/11/19. doi: 10.1016/j.pnpbp.2015.11.003. PubMed PMID: 26577297.

20. Bailey CP, Husbands SM. Novel approaches for the treatment of psychostimulant and opioid abuse – focus on opioid receptor-based therapies. Expert Opin Drug Discov. 2014;9(11):1333–44. Epub 2014/09/26. doi: 10.1517/17460441.2014.964203. PubMed PMID: 25253272; PubMed Central PMCID: PMCPMC4587358.

21. Jacobsen JH, Hutchinson MR, Mustafa S. Drug addiction: targeting dynamic neuroimmune receptor interactions as a potential therapeutic strategy. Curr Opin Pharmacol. 2016;26:131–7. Epub 2015/12/15. doi: 10.1016/j.coph.2015.10.010. PubMed PMID: 26657076.

22. Melik Parsadaniantz S, Rivat C, Rostene W, Reaux-Le Goazigo A. Opioid and chemokine receptor crosstalk: a promising target for pain therapy? Nat Rev Neurosci. 2015;16(2):69–78. Epub 2015/01/16. doi: 10.1038/nrn3858. PubMed PMID: 25588373.

23. Nutt DJ, Lingford-Hughes A, Erritzoe D, Stokes PR. The dopamine theory of addiction: 40 years of highs and lows. Nat Rev Neurosci. 2015;16(5):305–12. Epub 2015/04/16. doi: 10.1038/nrn3939. PubMed PMID: 25873042.

24. Piechota M, Korostynski M, Solecki W, Gieryk A, Slezak M, Bilecki W, et al. The dissection of transcriptional modules regulated by various drugs of abuse in the mouse striatum. Genome Biol. 2010;11(5):R48. Epub 2010/05/13. doi: 10.1186/gb-2010-11-5-r48. PubMed PMID: 20459597; PubMed Central PMCID: PMCPMC2898085.

25. Tanga FY, Raghavendra V, DeLeo JA. Quantitative real-time RT-PCR assessment of spinal microglial and astrocytic activation markers in a rat model of neuropathic pain. Neurochem Int. 2004;45(2-3):397–407. Epub 2004/05/18. doi: 10.1016/j.neuint.2003.06.002. PubMed PMID: 15145554.

26. Tanga FY, Nutile-McMenemy N, DeLeo JA. qThe CNS role of Toll-like receptor 4 in innate neuroimmunity and painful neuropathy. Proc Natl Acad Sci U S A. 2005;102(16):5856–61. Epub 2005/04/06. doi: 10.1073/pnas.0501634102. PubMed PMID: 15809417; PubMed Central PMCID: PMCPMC556308.

27. Vabulas RM, Ahmad-Nejad P, Ghose S, Kirschning CJ, Issels RD, Wagner H. HSP70 as endogenous stimulus of the Toll/interleukin-1 receptor signal pathway. J Biol Chem. 2002;277(17):15107–12. Epub 2002/02/14. doi: 10.1074/jbc.M111204200. PubMed PMID: 11842086.

28. Tsan MF, Gao B. Cytokine function of heat shock proteins. Am J Physiol Cell Physiol. 2004;286(4):C739–44. Epub 2004/03/06. doi: 10.1152/ajpcell.00364.2003. PubMed PMID: 15001423.

29. Ndengele MM, Cuzzocrea S, Masini E, Vinci MC, Esposito E, Muscoli C, et al. Spinal ceramide modulates the development of morphine antinociceptive tolerance via peroxynitrite-mediated nitroxidative stress and neuroimmune activation. J Pharmacol Exp Ther. 2009;329(1):64–75. Epub 2008/11/27. doi: 10.1124/jpet.108.146290. PubMed PMID: 19033555; PubMed Central PMCID: PMCPMC2670603.

30. Buchanan MM, Hutchinson M, Watkins LR, Yin H. Toll-like receptor 4 in CNS pathologies.J Neurochem. 2010;114(1):13–27. Epub 2010/04/21. doi: 10.1111/j.1471-4159.2010.06736.x. PubMed PMID: 20402965; PubMed Central PMCID: PMCPMC2909662.

31. Perry VH, Hume DA, Gordon S. Immunohistochemical localization of macrophages and microglia in the adult and developing mouse brain. Neuroscience. 1985;15(2):313–26. Epub 1985/06/01. PubMed PMID: 3895031.

32. Grace PM, Hutchinson MR, Maier SF, Watkins LR. Pathological pain and the neuroimmune interface. Nat Rev Immunol. 2014;14(4):217–31. Epub 2014/03/01. doi: 10.1038/nri3621. PubMed PMID: 24577438; PubMed Central PMCID: PMCPMC5525062.

33. Guegan T, Cebria JP, Maldonado R, Martin M. Morphine-induced locomotor sensitization produces structural plasticity in the mesocorticolimbic system dependent on CB1-R activity. Addict Biol. 2016;21(6):1113–26. Epub 2016/10/26. doi: 10.1111/adb.12281. PubMed PMID: 26179931.

34. Suarez-Roca H, Abdullah L, Zuniga J, Madison S, Maixner W. Multiphasic effect of morphine on the release of substance P from rat trigeminal nucleus slices. Brain Res. 1992;579(2):187–94. Epub 1992/05/08. PubMed PMID: 1378346.

35. Glaum SR, Miller RJ, Hammond DL. Inhibitory actions of delta 1-, delta 2-, and mu-opioid receptor agonists on excitatory transmission in lamina II neurons of adult rat spinal cord. J Neurosci. 1994;14(8):4965–71. Epub 1994/08/01. PubMed PMID: 8046463.

36. Grudt TJ, Williams JT. mu-Opioid agonists inhibit spinal trigeminal substantia gelatinosa neurons in guinea pig and rat. J Neurosci. 1994;14(3 Pt 2):1646–54. Epub 1994/03/01. PubMed PMID: 8126561.

37. Stellwagen D, Beattie EC, Seo JY, Malenka RC. Differential regulation of AMPA receptor and GABA receptor trafficking by tumor necrosis factor-alpha. J Neurosci. 2005;25(12):3219–28. Epub 2005/03/25. doi: 10.1523/JNEUROSCI.4486-04.2005. PubMed PMID: 15788779.

38. Raghavendra V, Rutkowski MD, DeLeo JA. The role of spinal neuroimmune activation in morphine tolerance/hyperalgesia in neuropathic and sham-operated rats. J Neurosci. 2002;22(22):9980–9. Epub 2002/11/13. PubMed PMID: 12427855.

39. Zhang N, Rogers TJ, Caterina M, Oppenheim JJ. Proinflammatory chemokines, such as C-C chemokine ligand 3, desensitize mu-opioid receptors on dorsal root ganglia neurons. J Immunol. 2004;173(1):594–9. Epub 2004/06/24. PubMed PMID: 15210821.

40. Zhang RX, Li A, Liu B, Wang L, Ren K, Zhang H, et al. IL-1ra alleviates inflammatory hyperalgesia through preventing phosphorylation of NMDA receptor NR-1 subunit in rats. Pain. 2008;135(3):232–9. Epub 2007/08/11. doi: 10.1016/j.pain.2007.05.023. PubMed PMID: 17689191; PubMed Central PMCID: PMCPMC2323207.

41. Iwasaki A, Medzhitov R. Toll-like receptor control of the adaptive immune responses. Nat Immunol. 2004;5(10):987–95. Epub 2004/09/30. doi: 10.1038/ni1112. PubMed PMID: 15454922.

42. Hong J, Cho IH, Kwak KI, Suh EC, Seo J, Min HJ, et al. Microglial Toll-like receptor 2 contributes to kainic acid-induced glial activation and hippocampal neuronal cell death. J Biol Chem. 2010;285(50):39447–57. Epub 2010/10/07. doi: 10.1074/jbc.M110.132522. PubMed PMID: 20923777; PubMed Central PMCID: PMCPMC2998094.

43. Maze I, Covington HE, 3rd, Dietz DM, LaPlant Q, Renthal W, Russo SJ, et al. Essential role of the histone methyltransferase G9a in cocaine-induced plasticity. Science. 2010;327(5962):213–6. Epub 2010/01/09. doi: 10.1126/science.1179438. PubMed PMID: 20056891; PubMed Central PMCID: PMCPMC2820240.

44. Auerbach RK, Chen B, Butte AJ. Relating genes to function: identifying enriched transcription factors using the ENCODE ChIP-Seq significance tool. Bioinformatics. 2013;29(15):1922–4. Epub 2013/06/05. doi: 10.1093/bioinformatics/btt316. PubMed PMID: 23732275; PubMed Central PMCID: PMCPMC3712221.

45. Blattler A, Yao L, Wang Y, Ye Z, Jin VX, Farnham PJ. ZBTB33 binds unmethylated regions of the genome associated with actively expressed genes. Epigenetics Chromatin. 2013;6(1):13. Epub 2013/05/23. doi: 10.1186/1756-8935-6-13. PubMed PMID: 23693142; PubMed Central PMCID: PMCPMC3663758.

46. Mulligan MK, Wang X, Adler AL, Mozhui K, Lu L, Williams RW. Complex control of GABA(A) receptor subunit mRNA expression: variation, covariation, and genetic regulation. PLoS One. 2012;7(4):e34586. Epub 2012/04/17. doi: 10.1371/journal.pone.0034586. PubMed PMID: 22506031; PubMed Central PMCID: PMCPMC3323555.

47. Godino A, Jayanthi S, Cadet JL. Epigenetic landscape of amphetamine and methamphetamine addiction in rodents. Epigenetics. 2015;10(7):574–80. Epub 2015/05/30. doi: 10.1080/15592294.2015.1055441. PubMed PMID: 26023847; PubMed Central PMCID: PMCPMC4622560.

48. Popovic R, Martinez-Garcia E, Giannopoulou EG, Zhang Q, Zhang Q, Ezponda T, et al. Histone methyltransferase MMSET/NSD2 alters EZH2 binding and reprograms the myeloma epigenome through global and focal changes in H3K36 and H3K27 methylation. PLoS Genet. 2014;10(9):e1004566. Epub 2014/09/05. doi: 10.1371/journal.pgen.1004566. PubMed PMID: 25188243; PubMed Central PMCID: PMCPMC4154646.

49. Cao R, Zhang Y. SUZ12 is required for both the histone methyltransferase activity and the silencing function of the EED-EZH2 complex. Mol Cell. 2004;15(1):57–67. Epub 2004/07/01. doi: 10.1016/j.molcel.2004.06.020. PubMed PMID: 15225548.

50. Schoofs T, Rohde C, Hebestreit K, Klein HU, Gollner S, Schulze I, et al. DNA methylation changes are a late event in acute promyelocytic leukemia and coincide with loss of transcription factor binding. Blood. 2013;121(1):178–87. Epub 2012/11/16. doi: 10.1182/blood-2012-08-448860. PubMed PMID: 23152544.

51. Higgins GA, Allyn-Feuer A, Athey BD. Epigenomic mapping and effect sizes of noncoding variants associated with psychotropic drug response. Pharmacogenomics. 2015;16(14):1565–83. Epub 2015/09/05. doi: 10.2217/pgs.15.105. PubMed PMID: 26340055.

52. Guelen L, Pagie L, Brasset E, Meuleman W, Faza MB, Talhout W, et al. Domain organization of human chromosomes revealed by mapping of nuclear lamina interactions. Nature. 2008;453(7197):948–51. Epub 2008/05/09. doi: 10.1038/nature06947. PubMed PMID: 18463634.

53. Asensio-Juan E, Gallego C, Martinez-Balbas MA. The histone demethylase PHF8 is essential for cytoskeleton dynamics. Nucleic Acids Res. 2012;40(19):9429–40. Epub 2012/08/02. doi: 10.1093/nar/gks716. PubMed PMID: 22850744; PubMed Central PMCID: PMCPMC3479184.

54. Loader C. Local regression and likelihood. New York: Springer; 1999. xiii, 290 p. p.

55. Wu W, Dave NB, Yu G, Strollo PJ, Kovkarova-Naumovski E, Ryter SW, et al. Network analysis of temporal effects of intermittent and sustained hypoxia on rat lungs. Physiol Genomics. 2008;36(1):24–34. Epub 2008/10/02. doi: 10.1152/physiolgenomics.00258.2007. PubMed PMID: 18826996; PubMed Central PMCID: PMCPMC2604785.

56. Sun JY, Loader CR. Simultaneous Confidence Bands for Linear-Regression and Smoothing. Ann Stat. 1994;22(3):1328–45. doi: DOI 10.1214/aos/1176325631. PubMed PMID: WOS:A1994QJ59900012.

57. Hartigan J, Wong M. A K-Means Clustering Algorithm. Journal of the Royal Statistical Society Series C (Applied Statistics). 1979;28(1):100–8.

58. Wu W, Bleecker E, Moore W, Busse WW, Castro M, Chung KF, et al. Unsupervised phenotyping of Severe Asthma Research Program participants using expanded lung data. J Allergy Clin Immun. 2014;133(5):1280–8. doi: 10.1016/j.jaci.2013.11.042. PubMed PMID: WOS:000335450700007.

59. Falcon S, Gentleman R. Using GOstats to test gene lists for GO term association. Bioinformatics. 2007;23(2):257–8. doi: 10.1093/bioinformatics/btl567. PubMed PMID: WOS:000243992400060.

60. Iorio F, Rittman T, Ge H, Menden M, Saez-Rodriguez J. Transcriptional data: a new gateway to drug repositioning? Drug discovery today. 2013;18(7):350–7.

61. Lamb J, Crawford ED, Peck D, Modell JW, Blat IC, Wrobel MJ, et al. The Connectivity Map: using gene-expression signatures to connect small molecules, genes, and disease. Science. 2006;313(5795):1929–35. doi: 10.1126/science.1132939. PubMed PMID: 17008526.

62. Lamb J. The Connectivity Map: a new tool for biomedical research. Nature reviews Cancer. 2007;7(1):54–60. doi: 10.1038/nrc2044. PubMed PMID: 17186018.

63. Kupershmidt I, Su QJ, Grewal A, Sundaresh S, Halperin I, Flynn J, et al. Ontology-based meta-analysis of global collections of high-throughput public data. PloS one. 2010;5(9):e13066.

